# Loss of erythroblasts in acute myeloid leukemia causes iron redistribution with clinical implications

**DOI:** 10.1101/2020.10.26.350116

**Authors:** Marta Lopes, Tiago L. Duarte, Maria J. Teles, Laura Mosteo, Sérgio Chacim, Eliana Aguiar, Joana Pereira-Reis, Mónica Oliveira, André M. N. Silva, Nuno Gonçalves, Gabriela Martins, Isabella Y Kong, Magnus Zethoven, Stephin Vervoort, Sandra Martins, Miguel Quintela, Edwin D Hawkins, Fernanda Trigo, João T Guimarães, José M Mariz, Graça Porto, Delfim Duarte

## Abstract

Acute myeloid leukemia (AML) is a heterogeneous disease with poor prognosis and limited treatment strategies. Determining the role of cell-extrinsic regulators of leukemic cells is vital to gain clinical insights into the biology of AML. Iron is a key extrinsic regulator of cancer but its systemic regulation remains poorly explored in AML. To address this question, we studied iron metabolism in AML patients at diagnosis and mechanisms involved using the syngeneic MLL-AF9-induced AML mouse model. We found that AML is a disorder with a unique iron profile not associated with inflammation or transfusion and characterized by high ferritin, low transferrin, high transferrin saturation (TSAT), and high hepcidin. The increased TSAT in particular, contrasts with observations in other cancer types and in anemia of inflammation. Using the MLL-AF9 mouse model of AML, we demonstrated that leukemic blasts take up iron and that the AML-induced loss of erythroblasts is responsible for iron redistribution and an increase in TSAT. We also show that elevated TSAT at diagnosis is independently associated with increased overall survival in AML and suggest that TSAT may be a relevant prognostic marker in AML.

## Introduction

Acute myeloid leukemia (AML) is a heterogeneous disease with increasing incidence. Despite significant advances in supportive therapy and the recent introduction of new drugs, the overall survival of AML patients remains low, particularly in older patients (1). The identification of specific genetic alterations has improved the classification and prognostic stratification of AML patients but the role of cell-extrinsic factors remains underexplored.

Leukemic cells depend on iron for DNA synthesis and proliferation yet iron metabolism remains ill-defined in AML. The link between iron and cancer is well recognized (2) and epidemiological studies have found an association between higher transferrin saturation (TSAT) and risk of cancer (3, 4). In AML, previous studies have proposed restricting iron availability through the use of iron chelators (5–7) and targeting CD71-expressing leukemic cells through anti-CD71 antibodies (7, 8). In contrast, a recent study showed that ferumoxytol, an iron oxide nanoparticle, has *in vivo* and *in vitro* anti-leukemic activity (9). In addition, iron has a significant role in shaping local microenvironments. Iron polarizes macrophages into a pro-inflammatory M1 subtype (10) and we have previously observed that the iron chelator desferoxamine (DFO) protects bone marrow (BM) endosteal blood vessels and non-malignant hematopoietic stem cells in AML-burdened mice (11). At the same time, DFO did not improve survival or reduce leukemia burden (11). Hepcidin, the key systemic iron regulator, is increased in AML patients undergoing hematopoietic stem cell transplantation (HSCT) (12), a clinical setting where analysis of iron metabolism is confounded by a history of red blood cell (RBC) transfusions. Altogether, there is a lack of understanding of if and how iron metabolism is altered by AML. We decided to address this question in the present study.

## Results and Discussion

We analyzed serum iron parameters of 84 AML patients at diagnosis (Supplemental Table 1) and observed a unique profile: high ferritin, low transferrin, normal to high serum iron and elevated TSAT (median 51.5%) (Figure 1A). The AML iron profile was not associated with a particular European Leukemia Net (ELN) risk subgroup (Supplemental Figure 1). Interestingly, early studies by Finch and others (13–15) also reported an exceptional phenotype of AML of low transferrin and increased TSAT consistent with our results, an observation that remained to be explained (16). Although AML is not an inflammatory disease, inflammation has been associated with myeloid malignancies (17) and ferritin is considered an acute phase protein. The observed AML-related iron phenotype is however clearly distinct from the profile associated with anemia of inflammation (AI) (Figure 1B, C), which is characterized by low serum iron and low TSAT, as shown in a group of patients with rheumatological disorders and anemia (Figure 1B; Supplemental Table 2). Whilst the altered iron kinetics in AML could, in principle, be explained by iron overload due to frequent RBC transfusions (18), we did not find an association between the iron profile of AML at diagnosis and a history of RBC transfusions (23 patients) (Figure 1D). The combination of low transferrin, high TSAT and high ferritin is observed in hemochromatosis. The *HFE* variant most commonly associated with hemochromatosis is however not overrepresented in AML (19), the iron status is comparable between AML patients with and without *HFE* variants (20) and we found high levels of hepcidin in AML patients at diagnosis (Figure 1E), which is not compatible with hemochromatosis.

**Figure 1 –.**
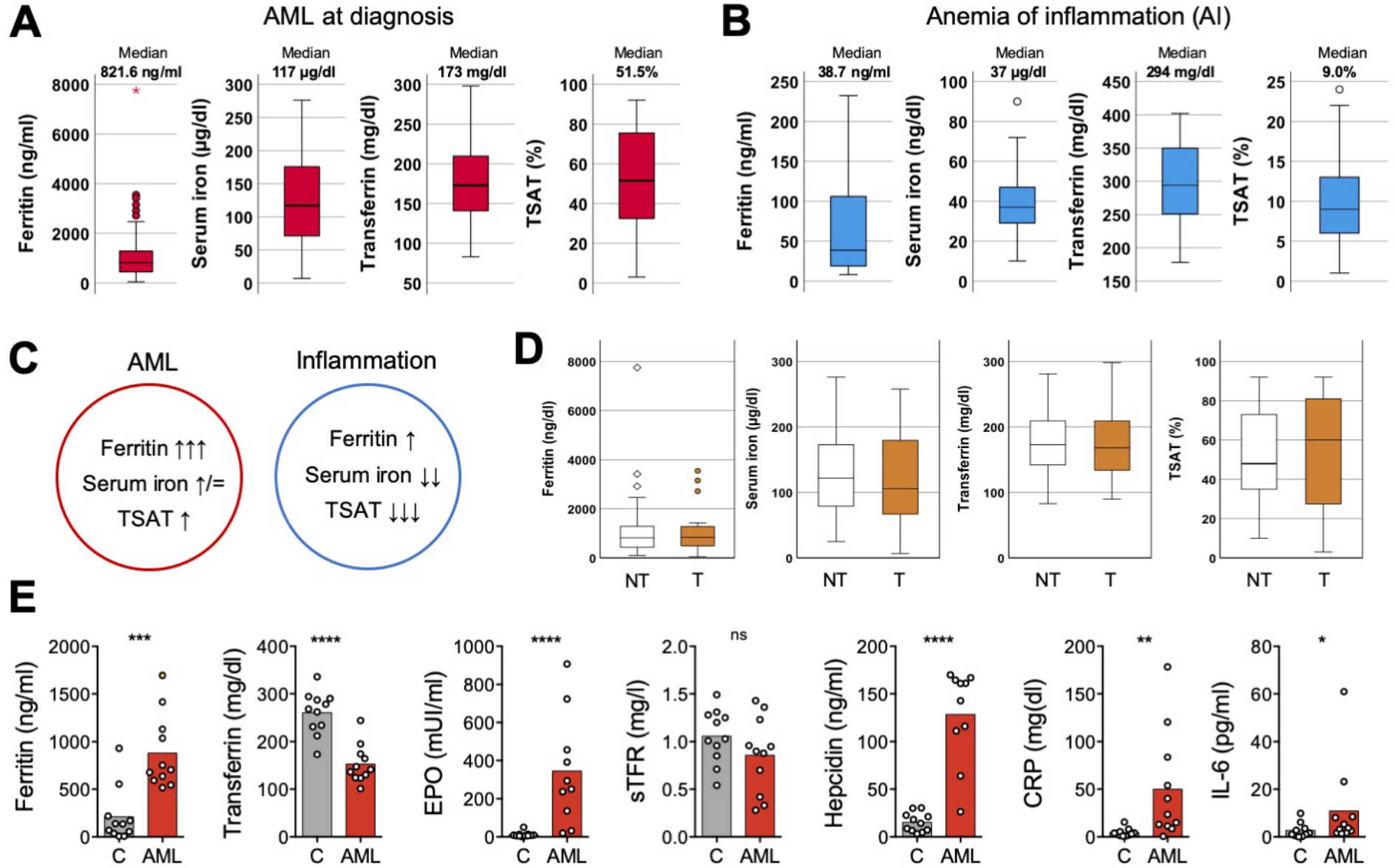
Iron metabolism in human AML at diagnosis. **(A)** Ferritin, serum iron, transferrin levels and transferrin saturation (TSAT) were analyzed in a cohort of 84 AML patients (Supplemental Table 1) at diagnosis. **(B)** An analagous evaluation was made in a group of 29 patients with rheumatological disorders (Supplemental Table 2) and with anemia of inflammation. **(C)** Summary of the key differences in iron profile observed in AML and in inflammation. **(D)** Comparison of the same parameters and in the same population described in (A), according to the transfusional status (NT - non transfused; T - transfused). **(E)** Iron-related parameters were analyzed in plasma collected from 11 patients with AML at diagnosis and from 12 patients with other conditions, mostly lymphoproliferative disorders (Supplemental Table 3).

To gain further insight into iron metabolism alterations in AML, we analyzed in more detail key iron-related parameters in 11 patients with newly diagnosed AML and 12 control patients with other hematological malignancies and inflammatory conditions (Supplemental Table 3). We found that AML patients had significantly higher ferritin (median 704.0 vs. 136.1 ng/ml; *p* = 0.003), lower transferrin (median 141.0 vs. 268.0 mg/dl; *p* < 0.001), higher erythropoietin (EPO) (272.0 vs. 8.3mUI/ml; *p* < 0.001) and very high circulating hepcidin levels (median 152.0 vs. 14.3 ng/ml; *p* < 0.001), albeit a modest increase in inflammatory C-reactive protein (CRP) and interleukin 6 (IL-6) (Figure 1E).

To explore the mechanism underlying the changes of iron metabolism in AML, we used the syngeneic MLL-AF9 AML mouse model. In this model, non-irradiated recipient mice are transplanted with AML cells expressing fluorescent proteins (EGFP or mTomato) or the congenic marker (CD45.1) that facilitates identification and discrimination of transformed cells from non-malignant counterparts by flow cytometry (Figure 2A) or imaging. MLL-AF9^+^ AML cells invade and expand primarily in the BM (Figure 2B) and spleen (Figure 2B, C), leading to hematopoietic failure. Typically, mice become fully infiltrated by leukemia from day 20 and succumb until day 35 if left untreated (Figure 2D). We analyzed these mice at clinically meaningful timepoints: at full infiltration and after treatment with induction chemotherapy (Figure 2E). The chemotherapy regimen was analogous to induction chemotherapy in patients and consisted of 5 days of cytarabine and 3 days of doxorubicin.

**Figure 2 –.**
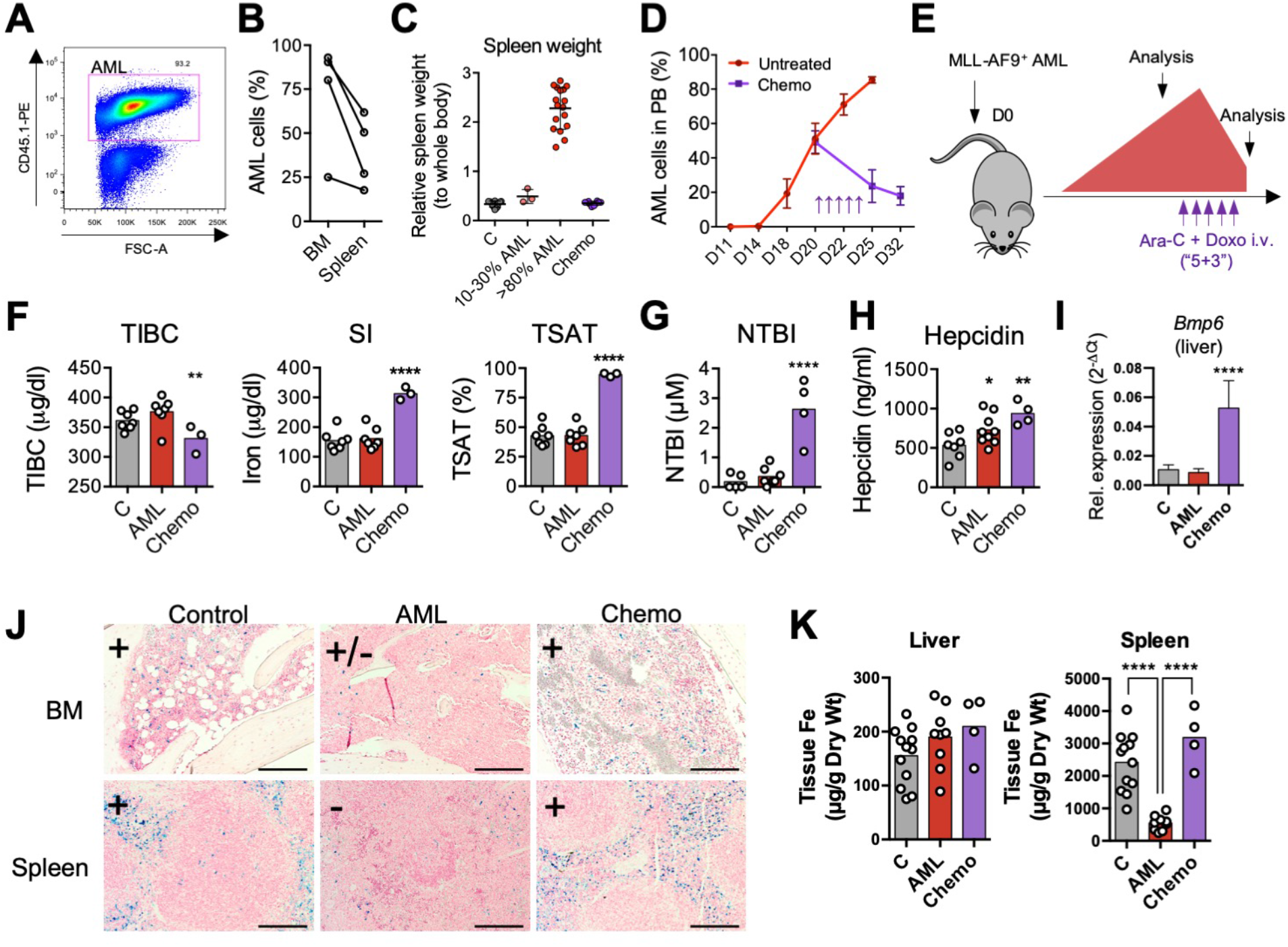
Iron metabolism in the MLL-AF9 AML mouse model. **(A)** MLL-AF9^+^ AML cells express a fluorescent protein or a specific marker, such as CD45.1 that allows the easy identification of AML cells by flow cytometry. **(B)** Mice transplanted with MLL-AF9^+^ AML cells are primarily infiltrated by leukemic blasts in the BM and in the spleen. n = 3 mice **(C)** The progressive infiltration of the spleen leads to increase weight that normalizes upon chemotherapy treatment. n = 12 (Control; C), 3 (10-30% AML), 17 (>80% AML) and 10 (post-chemotherapy; Chemo) mice. **(D)** Disease progression and response to chemotherapy (arrows; purple line) can be monitored by flow cytometry of the peripheral blood (PB). n = 5-20 mice per time point **(E)** Nonirradiated mice recipients were transplanted with 100,000 MLL-AF9^+^ AML cells and analyzed at full infiltration and after induction chemotherapy with cytarabine (Ara-C) and doxorubicin (Doxo). **(F)** Mice burden with AML had similar TIBC, SI, TSAT and NTBI levels. After chemotherapy, mice had increased SI levels, elevated TSAT and **(G)** presence of toxic NTBI. Each dot represents a mouse. **(H)** Circulating hepcidin levels were significantly increased in AML and after chemotherapy. **(I)** Hepatic *Bmp6* expression was significantly increased after chemotherapy treatment, as assessed by qPCR. n = 4-6 mice per group. **(J)** Representative images of Perls’ staining of BM and spleen sections from control, AML-burdened and chemotherapy-treated leukemic mice. Scale bar 200μm. **(K)** Quantification of non-heme iron levels in the liver and spleen. Each dot represents a mouse.

At full infiltration, no differences in total iron binding capacity (TIBC), serum iron or TSAT were detected (Figure 2F). In contrast, AML-burdened mice treated with chemotherapy had significantly lower TIBC, higher serum iron and very high TSAT (mean 94.6%). As expected, we detected toxic non-transferrin bound iron (NTBI) in sera after chemotherapy treatment (Figure 2G). This is consistent with the detection of NTBI after myeloablative chemotherapy (21) and HSCT (22) in clinical studies, which may contribute to tissue damage and infection. We also found higher levels of circulating hepcidin in mice infiltrated with AML and after chemotherapy (Figure 2H). We detected a marked increase in the hepatic expression of *Bmp6* in AML-infiltrated mice treated with chemotherapy (Figure 2I), suggesting that the increased circulating iron leads to increased hepcidin after chemotherapy.

We also analyzed iron deposition in three major sites of iron storage and recycling: the liver, spleen and BM. Perls’ staining revealed a loss of Prussian blue-positive cells in the BM and spleen of mice with AML, and a recovery after chemotherapy (Figure 2J). Quantification of non-heme iron revealed no differences in the liver but a decrease in the spleen iron content of AML-burdened mice (Figure 2K). These data show that in the mouse model, iron is not significantly deposited in the liver and is decreased in the BM and spleen, tissues that are infiltrated by AML cells.

The observation of increased TSAT in AML patients (Figure 1A) is not replicated in the mouse model (Figure 2F). A key difference between adult mice and humans is the significant contribution of the spleen to mouse erythropoiesis (23), both at steady state and upon acute hematopoietic stress. This contrasts with human extramedullary erythropoiesis, which is only significant under chronic BM stress, such as myelofibrosis. Consistently, AML-burdened mice had hyperleukocytosis with lymphopenia and thrombocytopenia but normal RBC and reticulocyte counts (Figure 3A), suggesting normal RBC output in AML. Analysis of the BM and spleen confirmed a depletion of BM erythroblasts (Figure 3B, C) but enhanced erythropoiesis in the spleen of leukemic mice (Figure 3D, E). Erythropoiesis in the liver was not observed (Supplemental Figure 2), showing that the spleen is the main site of erythropoiesis in mice infiltrated with AML. As expected, patients with AML (Supplemental Table 4) had decreased BM erythroblasts (Figure 3F, G; Supplemental Figure 3), consistent with an infiltrated BM that is unresponsive to EPO.

**Figure 3 –.**
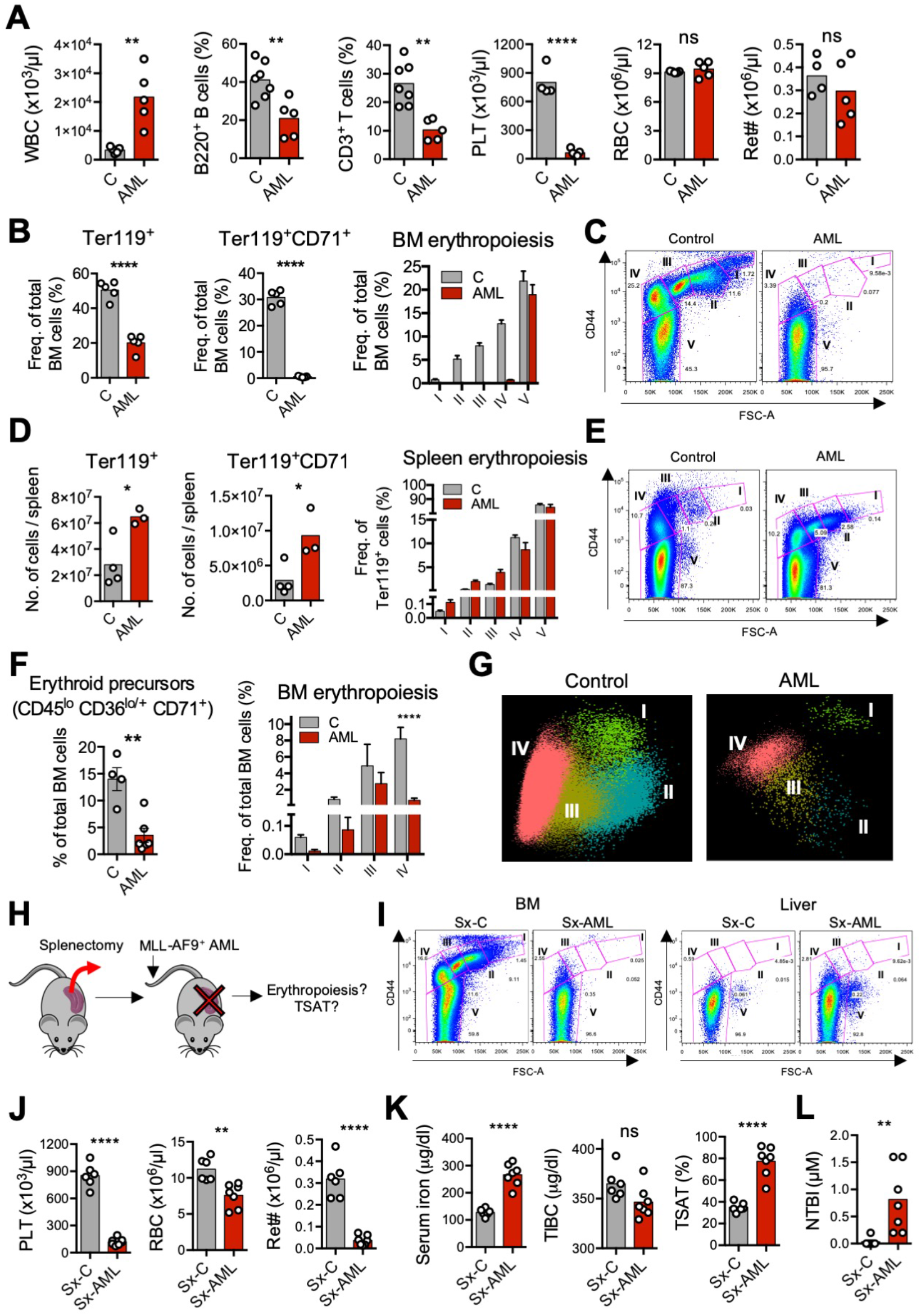
AML-induced loss of erythroblasts causes iron redistribution. **(A)** Peripheral blood counts for white blood cells (WBC), phenotypically-defined B and T cells, platelets (PLT), red blood cells (RBCs) and reticulocyte absolute numbers (Ret#). Each dot represents a mouse. **(B)** Flow cytometry analysis of BM Ter119^+^ and Ter119^+^CD71^+^ erythroid progenitors and of different erythropoiesis stages (I-V). n = 4 control and 3 AML mice. **(C)** Representative FACS plots of BM erythropoiesis in control and AML-burdened mice. **(D)** Flow cytometry analysis of spleen Ter119^+^ and Ter119^+^CD71^+^ erythroid progenitors and of different erythropoiesis stages (I-V). n = 4 control and 3 AML mice. **(E)** Representative FACS plots of BM erythropoiesis in control and AML-burdened mice. **(F)** Flow cytometry analysis of human BM erythroid precursors and of different erythropoiesis stages (I-IV). n = 4 control and 6 AML patients. **(G)** Representative Automatic Population Separator (APS) diagrams showing multiparametric analysis of flow cytometry data of human erythropoiesis in control and AML patients, based on the markers shown in Supplemental Figure 3. **(H)** Wild-type B6 mice were splenectomized (Sx), transplanted (Sx-AML) or not transplanted (Sx-C) with MLL-AF9^+^ AML and analyzed at full infiltration. **(I)** Representative FACS plots of BM and liver erythropoiesis in splenectomized mice. **(J)** Peripheral blood counts for PLT, RBCs and Ret# in splenectomized mice. **(K)** Sx-AML had increased SI and TSAT and **(L)** presence of toxic NTBI. Each dot represents a mouse.

Erythroblasts are the largest consumers of iron in the body, a requirement for hemoglobin synthesis. We hypothesized that the loss of erythroblasts in AML drives iron redistribution and TSAT increase, which is not observed in mice due to splenic erythropoiesis. We therefore predicted that splenectomized leukemic mice (Sx-AML) would phenocopy AML patients (Figure 3H). At full infiltration, BM erythroblasts were depleted and no erythropoietic compensation was observed in the liver of Sx-AML animals (Figure 3I). Analysis of the peripheral blood revealed thrombocytopenia, anemia and significant reticulocytopenia in Sx-AML mice (Figure 3J), consistent with loss of erythropoiesis. As predicted, Sx-AML mice had increased TSAT (Figure 3K) and detectable NTBI (Figure 3L), demonstrating that the loss of erythroblasts causes iron redistribution in AML.

We have shown that iron redistribution is partly corrected by chemotherapy (Figure 2J-K), which also leads to a dramatic increase in circulating iron (Figure 2F-G). This suggests that: 1) AML cells sequester a significant amount of the iron that is not being utilized for erythropoiesis, and 2) iron is released from dying cells after chemotherapy. Transferrin receptor 1 (CD71) is expressed at very high levels in erythroblasts but is not lineage-specific and is also expressed in proliferating cells, such as AML cells (Figure 4A). However, CD71 surface expression was lower in AML cells in comparison to proliferating Ki-67^hi^ non-malignant cells, suggesting decreased iron demand in leukemic cells (Figure 4B). Consistently, increased intracellular iron levels in AML cells were detected using electron microscopy and elemental analysis (EDS) (Figure 4C) quantification of the labile iron pool (LIP) with Phen Green SK labelling (24) using flow cytometry (Figure 4D). Furthermore, we detected increased expression of *Hmox1* and *Ftl1* in AML cells (Figure 4E), which is consistent with increased iron levels (25). In particular, a decreased heavy (*Fth1*) to light (*Ftl1*) ferritin chain ratio (Figure 4E) is indicative of increased LIP and is not altered in inflammation (26, 27). We also detected very low levels of FPN expression at the cell surface of AML cells (Figure 4F), an observation compatible with previous reports (9, 28) that may explain susceptibility to iron. Future studies should explore if the lower expression of FPN in leukemic blasts is the consequence of a disrupted oncogenic transcriptional program, of hepcidin-induced degradation or a combination of both.

**Figure 4 –.**
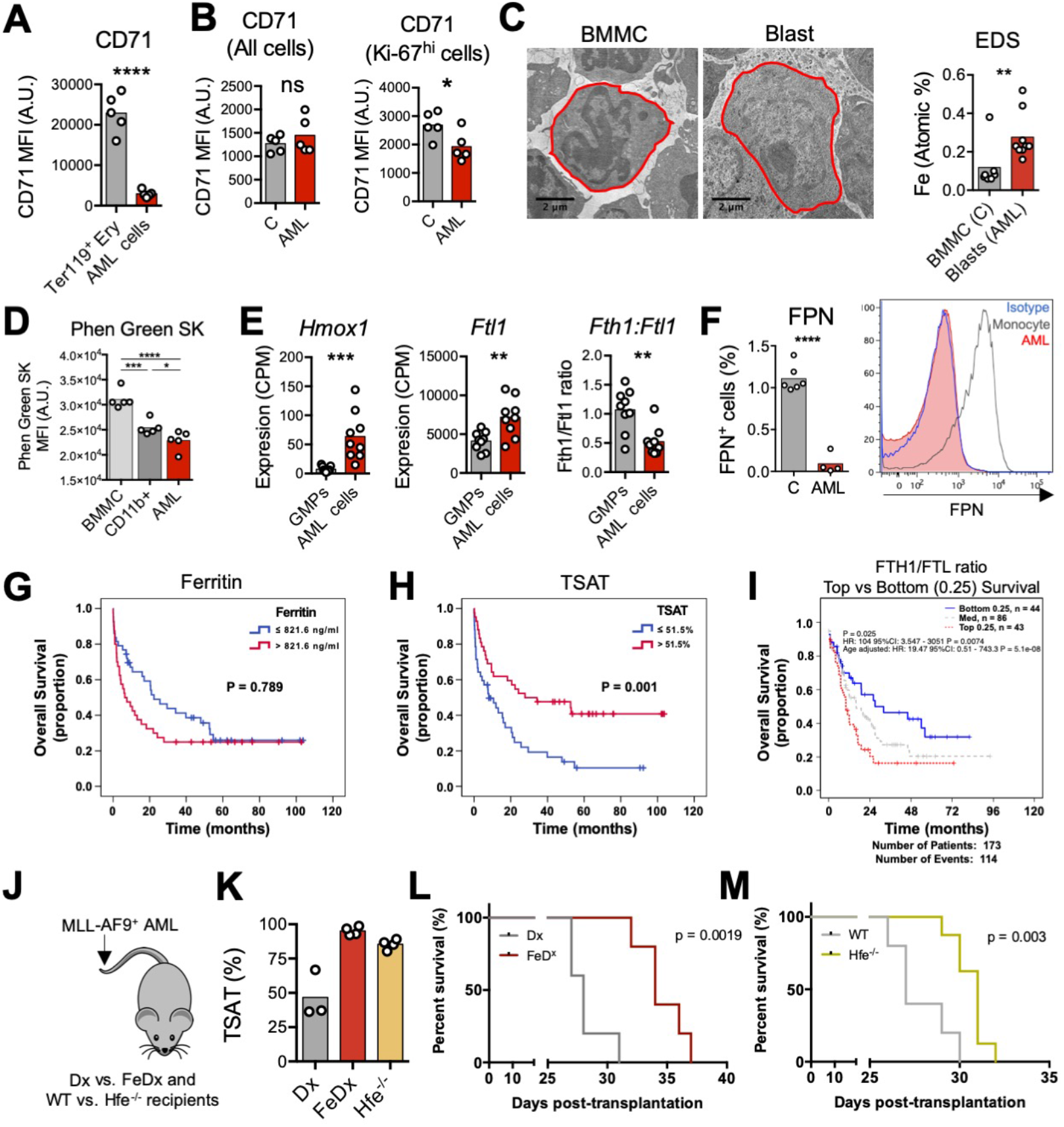
AML cells accumulate iron and increased TSAT at diagnosis is associated with increased OS. **(A)** FACS analysis of the mean fluorescence intensity (MFI) of surface CD71 in non-malignant Ter119^+^ erythroid precursors and AML blasts in the BM. n=5 mice per group. **(B)** Analysis of CD71 MFI in all or in proliferating Ki-67^hi^ control non-malignant BM cells and AML blasts isolated from the BM. n=5 mice per group. **(C)** Representative TEM images of BM cells from control and leukemic mice and EDS quantification of atomic iron (Fe) in selected cells. n=7-9 cells from 3 mice per group. **(D)** FACS analysis of LIP by loss of Phen Green SK intensity in BM mononuclear cells (BMMC) or CD11b^+^ cells from control mice or leukemic cells from AML-burdened mice. Each dot represents a mouse. Expression of **(E)** *Hmox1*, **(E)** *Ftl1* and ratio between *Fth1* and *Ftl1* expression in non-malignant granulocyte-macrophage progenitors (GMPs) and AML cells (GSE105159). Each dot represents a mouse. **(F)** Flow cytometry analysis of ferroportin (FPN) in non-malignant monocytes and AML cells. On the right: representative histogram. Each dot represents a mouse. Kaplan-Meier curves showing **(G)** no association between overall survival (OS) and ferritin levels (cutoff: median of 821 ng/ml), but **(H)** OS advantage for AML patients with elevated TSAT (cutoff: median of 51.5%). n=84 patients. **(I)** Kaplan-Meier curve showing improved OS of AML patients with lower heavy (FTH1) to light ferritin chain (FTL) expression ratio, a surrogate for increased LIP. Data from TCGA. **(J)** Mice transplanted with AML were treated with dextran (Dx) or iron-dextran (FeDx). On a separate cohort, WT or Hfe^-/-^ mice were transplanted with MLL-AF9^+^ AML. **(K)** FeDx-treated mice and Hfe^-/-^mice had increased TSAT. Kaplan-Meier curve showing improved survival of leukemic **(L)** FeDx and **(M)** Hfe^-/-^mice. n=5-8 mice per group.

The results above show that AML cells take up iron, although to a lower degree when compared to erythroblasts, and may be particularly susceptible to iron toxicity due to low FPN expression (9). This led us to hypothesize that increased circulating iron, as measured by TSAT, may have an impact on clinical outcomes in AML. We therefore analyzed our initial cohort of AML patients (Figure 1A) and observed that while ferritin levels were not associated with differences in overall survival (OS) (Figure 4G), AML patients with higher TSAT had a significantly better OS (Figure 4H). TSAT at diagnosis remained an independent prognostic factor for OS in a multivariate analysis including all patients (Supplementary Table 5) or only patients undergoing intensive chemotherapy treatment (Supplementary Table 6). Consistently, analysis of AML samples from The Cancer Genome Atlas (TCGA) revealed that lower FTH1 to FTL ratio, a surrogate of LIP (26, 27), was associated with improved OS (Figure 4I). To investigate whether increased TSAT prolongs survival in AML, we used two different mouse models of increased TSAT (Figure 4J, K). WT AML cells were transplanted into mice treated with iron-dextran (or dextran) or into mice deficient for the human homeostatic iron regulator protein (Hfe). We found that and survival was prolonged in mice treated with iron-dextran (Figure 4L) or in *Hfe*^-/-^ recipient mice (Figure 4M), suggesting that increased TSAT is not only associated with, but is rather involved in improving OS in AML. Future studies should explore the mechanism underlying this anti-leukemic effect, which may be direct, as suggested by the treatment with iron nanoparticles, indirect via the modulation of immune cells (such as macrophages). Furthermore, the prognostic role of TSAT in AML should be further explored in prospective studies.

In conclusion, we demonstrate that iron metabolism is dysregulated in AML (Supplemental Figure 4). Specifically, the three main findings of our study are that (1) AML is characterized by increased ferritin, low transferrin, high hepcidin and increased TSAT, (2) the loss of erythroblasts drives iron redistribution in AML, and (3) TSAT at diagnosis may be a useful prognostic marker to guide clinical decision in AML.

## Methods

Experimental procedures are provided in Supplemental Methods.

### Study approval

#### Human samples

Clinical and laboratorial data of patients with a diagnosis of AML (Supplemental Table 1) and AI (Supplemental Table 2) were retrospectively obtained after review and approval by Centro Hospital Universitário São João (CHUSJ)’s Local Ethics Committee (approval 220-20). Peripheral blood from patients with AML or lymphoproliferative diseases (Supplemental Table 3) were prospectively collected after informed consent. The study was reviewed and approved by Instituto Português de Oncologia do Porto (IPO-Porto)’s Local Ethics Committee (approval 158/018).

#### Mice

All animals received humane care according to the criteria outlined by the Federation of European Laboratory Animal Science Associations for the care and handling of laboratory animals (EU Directive 2010/63/EU). Experimental procedures were approved by i3S Animal Ethics Committee (DD_2019_15) and *Direção-Geral de Alimentação e Veterinária* (DGAV).

## Author Contributions

ML, TLD, LM, JR, MO and DD performed animal experiments; TLD performed qPCR experiments and tissue iron quantification; ML, MJT, NG, SM and JTG performed serum and plasma quantifications; SC, MQ, JMM and DD collected patient samples and performed analysis; EA and FT established a cohort of patients with iron quantifications and performed analysis; GM and DD performed FACS analysis of human samples; AS performed NTBI quantifications; IYK, MZ, SV and EDH performed RNA-seq and TCGA data analysis; DD and GP conceived the project and analyzed data; DD wrote the manuscript with contributions from all authors.

## Acknowledgments

The authors acknowledge the support of the i3S Scientific Platform HEMS, member of the national infrastructure PPBI - Portuguese Platform of Bioimaging (PPBI-POCI-01-0145-FEDER-022122). Funding for this project was provided by an EHA Research Grant award granted by the European Hematology Association to DD. This study was also supported by research funding from National Funds through FCT/MCTES – Portuguese Foundation for Science and Technology within the scope of the project UIDB/50006/2020 to AMNS and the Portuguese Society of Hematology, Fundação Amélia de Mello and National Blood Foundation to DD.

