## Supplemental for "Loss of erythroblasts in acute myeloid leukemia causes iron redistribution with clinical implications"

### SUPPLEMENTAL DATA

### Supplemental Methods

#### *Mice*

Animals were housed at the i3S animal facility under specific pathogen-free conditions, in a temperature and light-controlled environment, with free access to standard rodent chow (Teklad Global 14% Protein Rodent Maintenance Diet containing 175 mg/kg iron, Harlan Laboratories) and water. *Hfe*$^{-/-}$ mice were kindly donated by Helène Coppin (Toulouse, France) and backcrossed to C57BL/6 genetic background at the i3S animal house. C57BL/6J mice and CD45.1 were obtained from the i3S animal facility and from Charles River (France). Blood was obtained by retro-orbital bleeding under terminal anesthesia with isoflurane. All animals received humane care according to the criteria outlined by the Federation of European Laboratory Animal Science Associations for the care and handling of laboratory animals (EU Directive 2010/63/EU). Experimental procedures were approved by i3S Animal Ethics Committee (DD_2019_15) and Direção-Geral de Alimentação e Veterinária (DGAV).

#### *MLL-AF9 AML mouse model*

The model was established as described before (1, 2). For the generation of primary MLL-AF9-driven AML, granulocyte-monocyte progenitors (GMPs) were sorted from lineage-depleted BM cell suspensions obtained from WT, CD45.1 C57Bl/6 or mTmG donor mice using a FACS Aria II. GMPs were then transduced with retrovirus expressing the MLL-AF9 oncogene followed by IRES-GFP/dsRed and the transduction efficiency was determined by flow cytometry (using a FACS Canto II). pMIR/pMIG-FLAG-MLL-AF9 were a gift from Daisuke Nakada (Addgene plasmids # 71444 and 71443). 100-200k transduced GMPs were injected intravenously into sub-lethally irradiated (450 rads) WT C57Bl/6 recipient mice and primary AML blasts were harvested from these animals upon full disease infiltration (about 3 months post-transplantation).100,000 leukemic cells were injected into non-irradiated secondary and tertiary recipients.

#### *Mouse splenectomy*

The spleen of C57BL/6J 8-week female mice was removed under isoflurane anesthesia. A longitudinal incision was made on the left dorsolateral side of the abdomen, caudal to the last rib. The peritoneum was opened, the spleen was carefully mobilized using forceps. Splenic blood vessels were ligated, and the spleen was removed. The peritoneal cavity was closed with a PGA 5.0 continuous suture and the skin closed with PGA 5.0 individual interrupted suture. After surgery, the animals were allowed to recover for 9 days prior to MLL-AF9 AML transplantation. Mice were monitored closely and sacrificed at full infiltration.

***Drug treatments***

Induction chemotherapy was mimicked by treating young adults (8 to 10-week-old mice) infiltrated with AML (>day 20 post-transplant) with cytarabine (Ara-C; 100mg/kg/day I.V.) for 5 days and doxorubicin (Doxo; 3mg/kg/day I.V.) for 3 days. Ara-C and Doxo were obtained from IPO-Porto.

In the case of iron dextran treatment, mice received intraperitoneal (i.p.) injections of 1.25mg iron dextran (FeDx; Sigma, #D8517) or 1,7mg dextran control (Dx; Sigma, #D9260) every 4 days between days D7 and D27 after MLLAF9 cells transplantation (total of 6 injections of FeDX or Dx).

***Flow cytometry***

For mouse flow cytometry, mouse femurs and tibias were dissected and crushed in PBS containing 2% FBS. Bone marrow cells were transferred through a 40 µm cell strainer into a 50 ml tube and centrifuged at 500xg, 4°C for 5 min. Cells were incubated for 30 min at 4ºC with the following fluorochrome-conjugated antibodies: anti-CD71 APC (RI7217; 1:200), anti-Ter-119 APC/Cy7 (TER119; 1:200), anti-CD44 PE (IM7; 1:200), anti-Ly-6G/Ly-6C (Gr-1) PerCP (RB6-8C5; 1:200), anti-CD115 (CSF-1R) PE/Cy7 (AFS98; 1:200), anti-F4/80 BV510 (BM8; 1:100) all from Biolegend and anti-FPN AF647 (Novus Biologicals, cat # NBP1-21502AF647; 1:100). Flow cytometric quantification of the labile iron pool (LIP) was achieved with Phen Green™ SK diacetate (#P14313, Invitrogen™). Prior to use, Phen Green SK was dissolved in dimethylsulfoxide (DMSO). Bone marrow cells were incubated 5 min RT with Red Blood Cell Lysis Buffer (#420301, Biolegend), and centrifuged 500xg, 4°C for 5 min. Cells were incubated for 15 min at 37ºC with 5uM Phen Green™ SK diacetate diluted with Hank's balance salt solution (HBSS, Gibco, #14175). Staining for intracellular markers was performed using Foxp3 Transcription Factor Staining Buffer Set (Thermo Fisher Scientific, #00-5523-00). Isolated bone marrow cells were first stained for cell surface markers with the following fluorochrome-conjugated antibodies: anti-CD45.1 APC (A20, 1:100), and anti-CD71 FITC (RI7217, 1:200) all from Biolegend. Cells were then fixed and permeabilized with Fix:Perm solution for 30 min at RT. Cells were centrifuged at 500xg, 4ºC for 5 min and washed with the Permeabilization buffer. Cells were incubated with anti-Ki-67 Pe-Cy7 antibody (16A8, Biolegend) for 30 min at RT (3ul antibody diluted in 100ul of permeabilization buffer). Live and dead cells were distinguished using 4,6-diamidino-2-phenylindole (DAPI, Invitrogen). Calibrite Beads (BD Biosciences) were used to determine absolute cell counts (3). Cells were analyzed with a flow cytometer BD FACSCANTO II (BD Biosciences) and data were analyzed with FlowJo.

For human flow cytometry, BM aspirates were processed according to EuroFlow recommendations and using the following antibodies: anti-CD45 PacO (clone HI30) from Molecular Probes, anti-CD36 FITC (clone FA6.152), anti-CD117 APC or PE-Cy7 (clone 104D2D1) and anti-CD105 PE (clone IG2) from Beckman Coulter, anti-CD71 APC (clone M-A712) and anti-CD34 PerCP-Cy5.5 (clone 8G12) from BD Biosciences and HLA-DR PB (clone L243) from Biolegend. Cells were analyzed with a flow cytometer BD FACSCANTO II (BD Biosciences). Analysis of human BM erythropoiesis was performed as described in (4) using the software InfinicytTM (Cytognos S.L.).

***Hematologic and iron-related parameters***

Complete blood counts and erythrocyte indices were determined in a Sysmex XE-5000 (Emilio de Azevedo Campos) equipment in peripheral blood anticoagulated with EDTA, within less than 2 hours of collection. Serum or plasma samples and bone marrow fluid were obtained by centrifugation to determine erythropoietin, iron, transferrin and ferritin. The same sample was used also to determine soluble transferrin receptor and haptoglobin. Erythropoietin was determined using a chemiluminescent immunoassay on the Immulite 2000 analyzer (Siemens Healthcare Diagnostics); iron and transferrin were determined by a colorimetric method and ferritin by immunoturbidimetric assay, using Olympus reagents and analyzer (Olympus Diagnostics); soluble transferrin receptor and haptoglobin were determined by nephelometry techniques in the BNII nephelometer (Dade Behring). Internal and external controls and calibrators were used to ensure quality control in all the automatic analyzers, according to the local hospital laboratory procedures.

***Human IL-6 and hepcidin quantification***

IL-6 was analysed in plasma samples by a multiplexed immunoassay performed using a Human Premixed Multi-Analyte Kit (Magnetic Luminex Assay, Cat. # LXSAHM-04, Kit Lot Number L129889, R&D Systems, Inc., Minneapolis) and analyzed in the Luminex 200TM xMAPTM Technology (Luminex Corp., Netherlands). The results were quantified based on the Median Fluorescence Intensity (MFI) data using a standard five parameter logistic (5-PL) curve fit created by the Luminex xPONENT ® Software (version 3.1).

Plasma hepcidin levels in human patient samples were measured using a solid phase enzyme-linked immunosorbent (ELISA) kit (#EIA-5782; DRG Hepcidin 25 (bioactive) HS ELISA; GmbH, Germany) based on the principle of competitive binding. The absorbance was determined at 450 nm with the Synergy™ Mx Microplate Reader (Gen5). Assay sensitivity is between 0.153 ng/ml - 81 ng/ml.

***Mouse serum analysis***

Serum iron and total iron binding capacity (TIBC) were analyzed using a Beckman Coulter A5800 clinical chemistry analyzer.

Mouse serum hepcidin was measured using a Hepcidin-Murine Compete™ ELISA kit (#HMC-001, Intrinsic Lifesciences, La Jolla, CA, USA) based on the principle of competitive binding and designed for specific quantification of hepcidin-1 (hepc-1). The absorbance was determined at 450 nm with the Synergy™ Mx Microplate Reader (Gen5). Assay sensitivity is between 0.14 - 1000 ng/ml.

#### *Quantitative polymerase chain reaction (qPCR)*

Liver fragments were homogenized in NZYol (NZYTech, Lisbon, Portugal). The cell suspension was filtered through a 40 μm nylon strainer (BD Falcon) to a 50 ml tube and centrifuged at 500×g for 5 min at 4°C. Total RNA was extracted using NZYol (NZYTech, Lisbon, Portugal), followed by DNase treatment (Turbo DNA-free kit) (Life Technologies, Carlsblad, CA, USA). RNA integrity was assessed with an Experion Automated Electrophoresis System (Bio-Rad, Hercules, CA, USA). First-strand cDNA was prepared from 2 μg RNA using the NZY First-Strand cDNA Synthesis Kit (NZYTech, Lisbon, Portugal) with an oligo(dT)$_{18}$ primer. Relative gene expression levels were quantified using a CFX384 Real-Time PCR Detection System (Bio-Rad, Hercules, CA, USA). Primer sequences are listed in Supplementary Methods. All reactions were performed in a total volume of 10 μl with iTaq™ Universal SYBR Green Supermix. The amplification protocol consisted of denaturation at 95°C for 3 min 30 s and 40 cycles of 95°C for 20 s, and 59°C for 30 s. The relative quantity of each transcript was estimated by the $2^{-\Delta Ct}$ method, after normalization against the averaged Ct of two endogenous control genes, Hypoxanthine phosphoribosyltransferase 1 (Hprt1) and beta Actin (Actb).

Primer sequences:

| Gene ID | Forward primer sequence (5'-3') | Reverse primer sequence (5'-3') |
|---|---|---|
| Actb | GGCGGACTGTTACTGAGCTGCGTTT | AAAGCCATGCCAATGTTGTCTCTT |
| Bmp6 | TCCCCACATCAACGACACCA | TCCCCACCACACAGTCCTTG |
| Hprt1 | AGATGGGAGGCCATCACATTGT | ATGTCCCCCGTTGACTGATCAT |

#### *Perls staining*

Femurs were fixed in neutral formalin 10%, incubated in EDTA-based decalcifying solution for 48 h and embedded in paraffin. Spleen samples were fixed in neutral formalin 10% and embedded in paraffin. Following deparaffinization with xylene and hydration by a passage through a grade of alcohols, 3 μm-thick spleen and 4 μm-thick femur sections were stained with Perls' Prussian blue reaction for ferric iron at IPATIMUP Diagnostics. All slides were examined by the same observer (TLD), in a blind manner.

#### *Non-heme iron quantification*

Non-heme iron content in spleen and liver samples was measured by the bathophenantroline method as described before (5).

#### Non-transferrin bound iron (NTBI) determination

NTBI determinations were carried out by the ultracentrifugation assay first described by Singh et al.(6) Serum samples (90 μl) were treated with 10 μl of an 800 mM NTA solution at pH 7.0 and were allowed to incubate for 30 minutes at room temperature. The mixture was then ultra-filtered at 10000xg and 4ºC for 1 hour, using 10 kDa molecular weight cut-off Amicon Ultra 0.5 ml filtration units (Millipore). Iron content was subsequently measured in the ultra-filtrate by the colorimetric ferrozine assay adapted to a 96-microwell plate format(7, 8). 50 µl sample was mixed with 20 µl 4 mM ascorbic acid solution prepared in formic acid (0.2 M at pH 3) and allowed to incubate in the dark at room temperature. After 5 min, 25 µl 1 mM ferrozine reagent prepared in formic acid solution was added and the incubation was allowed to proceed for further 30 min. Iron concentration is read from a standard curve after determination of the solution absorbance at 562 nm. Standard iron solutions (0–10 µM) were prepared in 80 mM NTA and were analysed by using the same procedures as with serum samples.

***Electron microscopy and EDS***

The transmission electronic microscopy and energy-dispersive X-ray spectroscopy (EDS) analysis were performed at the HEMS core facility at i3S, University of Porto, Portugal with the assistance of Ana Rita Malheiro e Rui Fernandes and according to previously published methods (9). Femoral epiphysis samples were fixed by immersion in 2.5% glutaraldehyde and 2% paraformaldehyde in 0.1M sodium cacodylate buffer (pH 7.4) solution for 5 days and subsequently incubated in EDTA-based decalcifying solution (MoL-Decalcifier) for 48h. After washing and two hours in post-fixating 2% osmium tetroxide in 0.1M sodium cacodylate buffer (pH 7.4) solution, tissues were washed in buffer, incubated with 1% Uranyl acetate overnight, washed in buffer and dehydrated through a graded series of ethanol, and embedded in Epon (EMS). Semithin sections (approximately 0.5 µm thick), stained with toluidine blue and methylene blue, were used to search for areas of interest. Ultrathin sections were cut at 50 nm and prepared on a RMC Ultramicrotome (PowerTome, USA) using a diamond knife and recovered to 200 mesh copper grids, followed by double contrast method with 2% uranyl acetate and saturated lead citrate solution. Visualization was performed at 80 kV in a (JEOL JEM 1400 microscope, Japan) and digital images were acquired using a CCD digital camera Orious 1100 W (Tokyo, Japan). For Energy-dispersive X-ray spectroscopy (EDS) analysis, unstained sections were mounted on formvar/carbon film-coated mesh nickel grids and a beryllium holder (EM-21150, Jeol Ltd.) was used. An X-Max 80 mm2 (Oxford Instruments, Bucks, England) operated at 120 kV was coupled to the microscope.

***TCGA data analysis***

RSEM (RNA-Seq by Expectation Maximization) (10) scaled expression values for TCGA were downloaded from the GDAC Firehose website (11). Counts were normalised to transcripts-per-million (TPM) values with a pseudo-count of 2. Entrez gene IDs were mapped to HGNC gene symbols using the biomaRt R package (version 2.42.0) (12) and collapsed to unique values per gene symbol by selecting the most variable entrez ID among all samples for each gene symbol. Primary tumour samples were selected using the TCGAbiolinks R package (version 2.14.0) (13) and were matched with clinical information from the TCGA Pan-Cancer Clinical Data Resource (14). The ratio of TPM values between the FTH1 and FTL genes was used to fit Kaplan-Meier and Cox regression models for the TCGA LAML cohort using the survival R package (version 3.1-8) (15).

***RNA-seq data analysis***

Transcriptome analysis were performed on previously published data GSE105159 (1). FeatureCounts (version 1.34.7) was used for read counting after which differential gene expression analysis was performed using Voom-LIMMA packages (limma version 3.40.6)(16-18).

***Statistical analysis***

Data were analyzed using GraphPad Prism (GraphPad Software, CA, USA) and SPSS Statistics (IBM). Unpaired t-test was used to compare group means. One-way ANOVA with post hoc Tukey test or Bonferroni correction was used for multiple comparisons. For survival analysis, the log-rank test was performed. Differences were considered significant when $p < 0.05$ (*$p < 0.05$, **$p < 0.01$, ***$p < 0.001$).

**References (methods)**

11. Center BITGDA. Analysis-ready standardized TCGA data from Broad GDAC Firehose 2016_01_28 run. 2016.
12. Durinck S, Spellman PT, Birney E, and Huber W. Mapping identifiers for the integration of genomic datasets with the R/Bioconductor package biomaRt. *Nature protocols.* 2009;4(8):1184.
13. Colaprico A, Silva TC, Olsen C, Garofano L, Cava C, Garolini D, et al. TCGAbiolinks: an R/Bioconductor package for integrative analysis of TCGA data. *Nucleic acids research.* 2016;44(8):e71-e.
14. Liu J, Lichtenberg T, Hoadley KA, Poisson LM, Lazar AJ, Cherniack AD, et al. An Integrated TCGA Pan-Cancer Clinical Data Resource to Drive High-Quality Survival Outcome Analytics. *Cell.* 2018;173(2):400-16 e11.
15. Therneau T. 2015.
16. Liao Y, Smyth GK, and Shi W. featureCounts: an efficient general purpose program for assigning sequence reads to genomic features. *Bioinformatics.* 2014;30(7):923-30.
17. Liao Y, Smyth GK, and Shi W. The R package Rsubread is easier, faster, cheaper and better for alignment and quantification of RNA sequencing reads. *Nucleic acids research.* 2019;47(8):e47-e.
18. Law CW, Chen Y, Shi W, and Smyth GK. voom: Precision weights unlock linear model analysis tools for RNA-seq read counts. *Genome biology.* 2014;15(2):R29.

### SUPLEMENTAL FIGURES

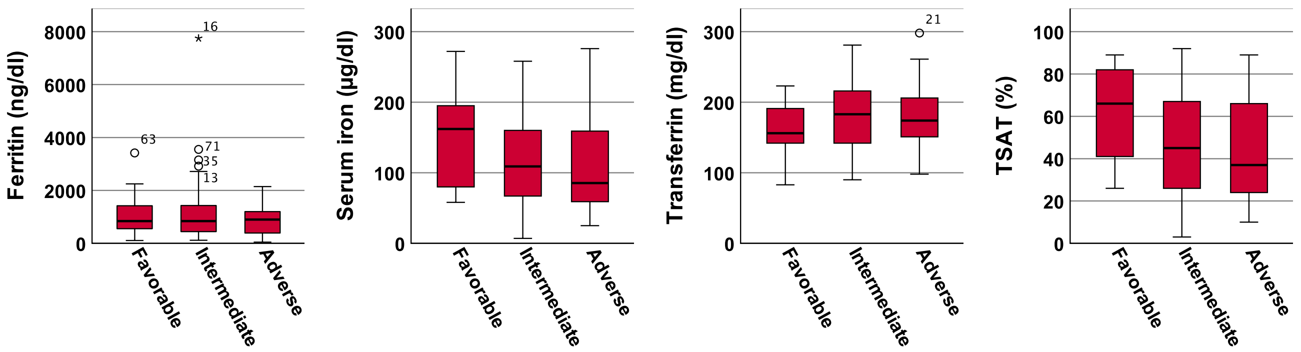


**Supplemental Figure 1.** Comparison of levels of ferritin, iron, transferrin and transferrin saturation (TSAT) in the sera of 84 AML patients (Supplemental Table 1), according to ELN risk subgroup (favorable, intermediate, adverse). No significance differences were detected.

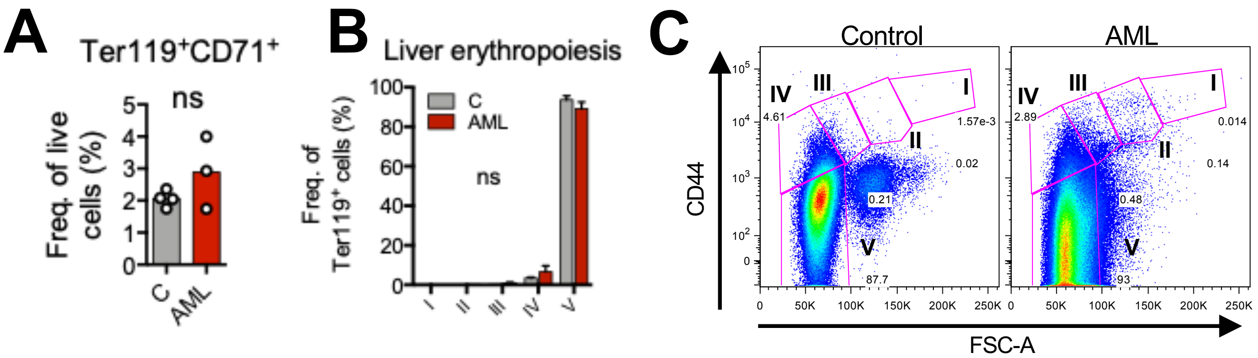


**Supplemental Figure 2. Analysis of erythropoiesis in the liver. (A)** Analysis of Ter119+CD71+ erythroid progenitors and **(B)** erythropoiesis stages revealed no significant differences between control and AML-burden mice. **(C)** Representative FACS plots of erythropoiesis stages in the liver. Data representative of 4 control and 3 leukemic mice.

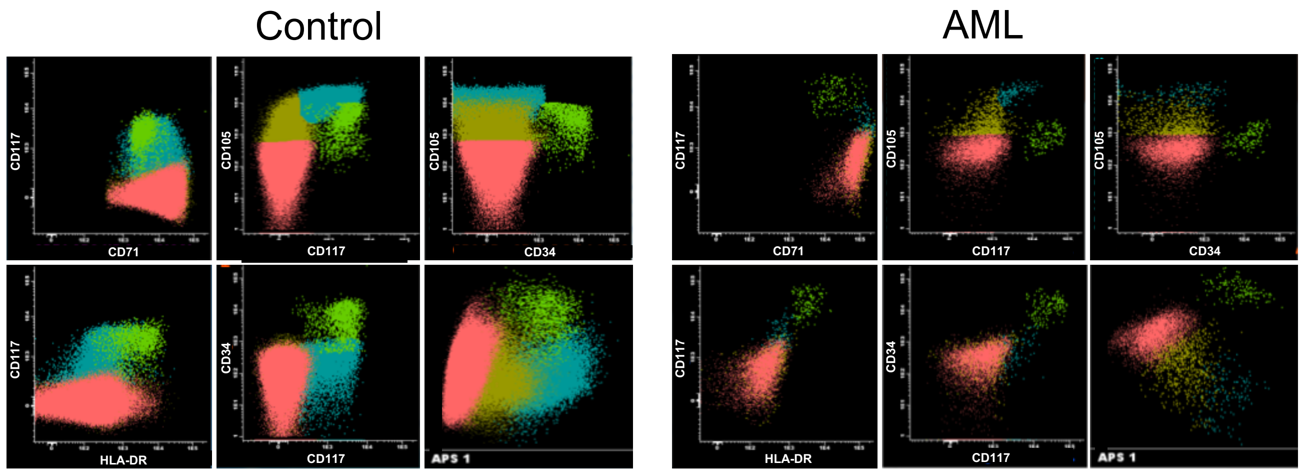


**Supplemental Figure 3. Analysis of human bone marrow erythropoiesis in patients without (control) and with AML.** Representative FACS plots showing different stages of human erythropoiesis. Ligh green dots: stage I nucleated red blood cells (NRBCs); blue dots: stage II NRBCs; dark green dots: stage III NRBCs; pink dots: stage IV NRBCs.

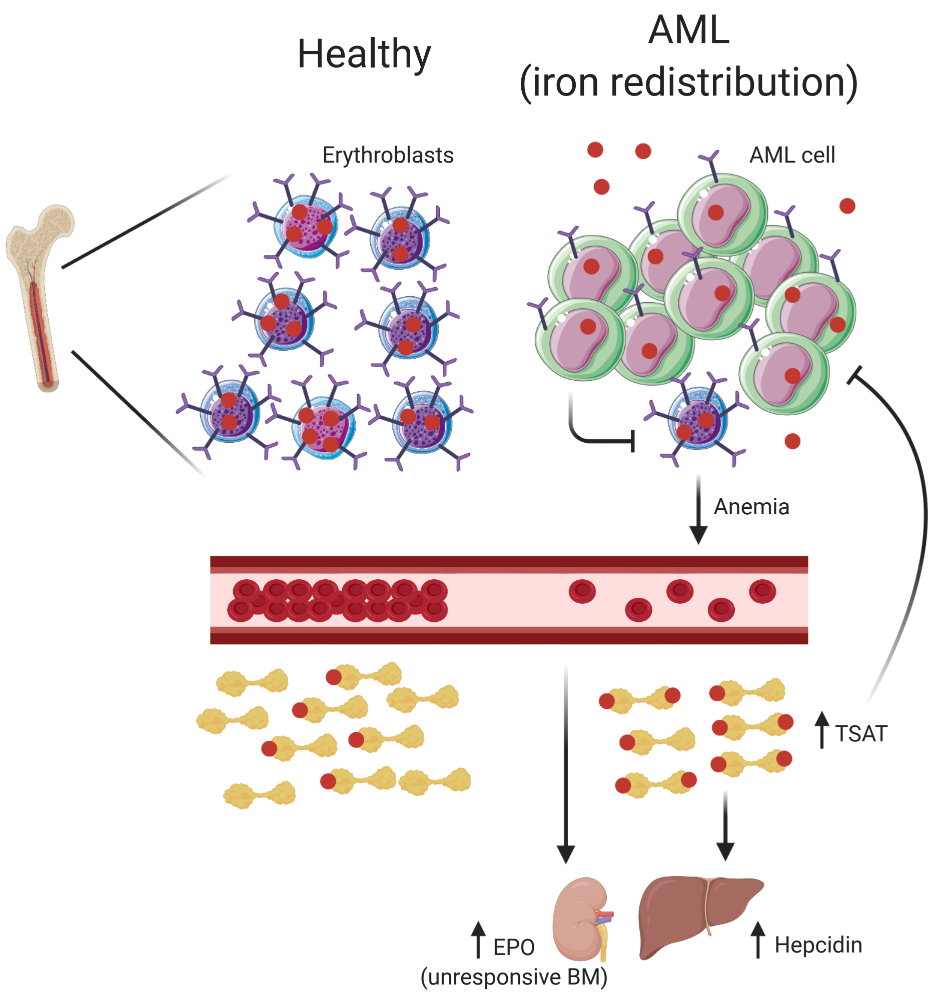


**Supplemental Figure 4.** In contrast with healthy individuals, patients with AML have proliferation of leukemic cells in the bone marrow (BM) and consequently loss of CD71$^{hi}$ erythroblasts, which causes iron redistribution and increased transferrin saturation (TSAT). In parallel to increased hepcidin, there are increased levels of erythropoietin (EPO) due to anemia but the BM is unresponsive. Increased TSAT is also associated with better overall survival in AML. Created with BioRender.com

### SUPPLEMENTAL TABLES

**Supplemental Table 1.** Baseline characteristics of 84 AML patients at diagnosis

| **Characteristics** | |
|---|---|
| **Gender, N (%)** | |
| Male | 48 (57,1) |
| Female | 36 (42,9) |
| **Median age, years (IQR)** | 62,5 (48,0-72,0) |
| **Median Hb, g/dl (IQR)** | 8,7 (7,5-9,9) |
| **Median WBC count, x10$^9$/l (IQR)** | 20,855 (7,260-45,558) |
| **Median platelet count, x10$^9$/l (IQR)** | 42,00 (26,50-94,75) |
| **Ferritin, ng/ml (IQR)** | 821,6 (445,1-1301,1) |
| **Serum iron, µg/dl (IQR)** | 117 (70,0-175,8) |
| **Transferrin, mg/dl (IQR)** | 173,0 (140,5-209,8) |
| **TSAT, % (IQR)** | 51,5 (31,8-76,8) |
| **Previous RBC transfusion, N (%)** | 23 (27,4) |
| **Therapy, N (%)** | |
| Intensive chemotherapy | 54 (64,3) |
| Hypomethylating agent | 6 (7,1) |
| Best supportive care | 22 (26,2) |
| **CR/CRi, N (%)** | 50 (59,4) |
| **Cytogenetic Risk, N (%)** | |
| Favorable | 16 (19,0) |
| Intermediate | 56 (66,7) |
| Adverse | 10 (11,9) |
| **AML subtype, N (%)** | |
| *de novo* AML | 70 (83,3) |
| Secondary AML | 14 (16,7) |
| **WHO classification 2016, N (%)** | |
| AML with recurrent genetic abnormalities | |
| AML with t(8;21)(q22;q22.1); RUNX1-RUNX1T1 | 3 (3,6) |
| AML with inv(16)(p13.1q22) or t(16;16)(p13.1;q22); CBFB-MYH11 | 7 (8,3) |
| AML with t(9;11)(p21.3;q23.3);MLLT3-KMT2A | 1 (1,2) |
| AML with inv(3)(q21.3q26.2) or t(3;3)(q21.3;q26.2); GATA2, MECOM | 2 (2,4) |
| Provisional entity: AML with BCR-ABL1 | 1 (1,2) |
| AML with mutated NPM1 | 22 (26,2) |
| AML with myelodysplasia-related changes | 21 (25,0) |
| Therapy-related myeloid neoplasms | 2 (2,4) |
| AML, NOS | |
| AML without maturation | 3 (3,6) |
| AML with maturation | 6 (7,1) |
| Acute myelomonocytic leukemia | 4 (4,8) |
| Acute monoblastic/monocytic leukemia | 6 (7,1) |
| Pure erythroid leukemia | 1 (1,2) |
| Not classified | 5 (6,0) |

N - number of observations; IQR - interquartile range; Hb - hemoglobin; WBC - peripheral white blood cell count; CR - complete response; CRi - complete response with incomplete blood count recovery.

**Supplemental Table 2.** Baseline characteristics of 29 rheumatological patients with anemia of inflammation

| **Characteristics** | |
|---|---|
| **Gender, N (%)** | |
| Male | 9 (31) |
| Female | 20 (69) |
| **Median age, years (IQR)** | 57,0 (48,5-72,0) |
| **Median Hb, g/dl (IQR)** | 11,4 (10,9-11,9) |
| **Median WBC count, x10$^9$/l (IQR)** | 7,240 (5,260-9,650) |
| **Median platelet count, x10$^9$/l (IQR)** | 258,00 (210,50-318,25) |
| **Ferritin, ng/ml (IQR)** | 38,7 (17,8-119,9) |
| **Serum iron, µg/dl (IQR)** | 37,0 (28,0-49,0) |
| **Transferrin, mg/dl (IQR)** | 294,0 (244,0-355,5) |
| **TSAT, % (IQR)** | 9,0 (6,0-13,0) |
| **Oral iron treatment, N (%)** | 9 (31,0) |
| **Diagnosis, N (%)** | |
| Juvenile idiopathic arthritis | 1 (3,4) |
| Rheumatoid arthritis | 15 (51,7) |
| Psoriatic arthritis | 5 (17,2) |
| Systemic lupus erythematosus | 4 (13,8) |
| Behçet's disease | 1 (3,4) |
| Ankylosing spondylitis | 2 (6,9) |
| Sjögren syndrome | 1 (3,4) |

N - number of observations; IQR - interquartile range; Hb - hemoglobin; WBC - peripheral white blood cell count.

**Supplemental Table 3.** Characteristics of hematological patients at diagnosis

| Group | ID | Diagnosis | Age | Sex | Cytogenetics | Molecular Genetics |
|---|---|---|---|---|---|---|
| Control | DC07 | Folicular Lymphoma | 44 | F | | |
| | DC10 | B ALL in remission | 63 | F | | |
| | DC14 | NHL MALT | 50 | M | | |
| | DC16 | Folicular Lymphoma | 51 | M | | |
| | DC22 | NHL MALT | 78 | F | | |
| | DC23 | NHL DLBCL | 45 | M | | |
| | DC24 | MCL | 83 | M | | |
| | DC25 | NHL MALT | 45 | F | | |
| | DC28 | Sarcoidosis | 59 | F | | |
| | DC29 | NHL MALT | 51 | F | | |
| | DC30 | Folicular Lymphoma | 70 | F | | |
| | DC31 | NHL MALT | 54 | M | | |
| AML | DC01 | De novo AML | 69 | F | 46,XY | IDH1/2- |
| | DC02 | De novo AML | 65 | F | 47,XX,+8[2]/47,idem,t(3;21)(p25;q22)[18] | FLT3-ITD+ |
| | DC03 | De novo AML | 22 | M | 46,XY,inv(16)(p13q22)[20] | CBFB-MYH11+/ FLT3-ITD+ |
| | DC04 | De novo AML | 64 | F | No metaphases | |
| | DC06 | De novo AML | 57 | F | 46,XX[20] | FLT3-ITD- |
| | DC08 | De novo AML | 69 | F | 46,XX[8] | FLT3-ITD- |
| | DC17 | AML secundary to MDS/MPN | 62 | F | 46,XX,add(3)(q12),add(18)(q23)x2[6] | FLT3-ITD- |
| | DC18 | De novo AML | 53 | M | 44~46,XY,add(11)(q23),inc[4] | |
| | DC20 | De novo AML | 48 | M | 45,X,-Y,t(8;21)(q22;q22),del(9)(q21q34)[20] | RUNX1-RUNX1T1+ |
| | DC26 | De novo AML | 57 | M | 46,XY[20] | |
| | DC32 | De novo AML | 60 | M | 45~46,XY,i(1)(p10),-4,add(4)(p12),-5,add(7)(q32),+8,add(14)(p11.2),-15,-16,-17,i(21)(q10),+4mar[cp18]/46,XY[2] | |

ALL – acute lymphoblastic leukemia; NHL – non-Hodgkin lymphoma; MALT – mucosa-associated lymphoid tissue; MCL – mantle cell lymphoma; MDS – myelodysplastic syndrome; MPN – myeloproliferative neoplasm.

**Supplemental Table 4.** Characteristics of patients who underwent analysis of bone marrow nucleated red blood cells

| Group | Diagnosis | Age | Sex |
|---|---|---|---|
| Control | Multiple myeloma - 2 months after autologous HSC transplantation | 61 | M |
| | Anemia in characterization | 76 | F |
| | Allogeneic HSC transplantation post AML | 58 | M |
| | Relapse of Diffuse Large B-Cell Lymphoma | 81 | M |
| AML | De novo AML with 5q- | 48 | M |
| | De novo AML with normal karyotype | 84 | M |
| | De novo AML with trisomy 8 | 54 | F |
| | De novo AML with t(15;17) | 72 | M |
| | De novo AMKL with der(9)t(1;9) | 49 | F |
| | De novo AML with normal karyotype | 58 | F |

HSC – hematopoietic stem cell; AMKL – acute megakaryoblastic leukemia.

**Supplemental Table 5.** Multivariate Cox regression on overall survival of all 84 AML patients (Supplemental Table 1)

| | Overall Survival | | |
|---|---|---|---|
| **Variable** | HR | 95% CI | P |
| **Age** (>60 vs. ≤60 years) | 4,729 | 2,044-10,941 | **<0,001** |
| **ELN 2017** (intermediate vs. favorable risk) | 0,950 | 0,347-2,602 | 0,921 |
| **ELN 2017** (favorable vs. adverse risk) | 2,198 | 0,636-7,605 | 0,213 |
| **AML subtype** (de novo vs. secondary) | 0,663 | 0,212-2,071 | 0,479 |
| **TSAT** (≤51,5 vs. >51,5%) | 2,779 | 1,223-6,312 | **0,015** |
| **Response to induction** (CR vs. no CR) | 1,992 | 0,742-5,344 | 0,171 |

ELN – European Leukemia Net; TSAT – transferrin saturation; CR – complete response; HR – hazard ratio; CI – confidence interval; Bold: significant P values ($p<0.05$)

**Supplemental Table 6.** Multivariate Cox regression on overall survival of the AML patients who underwent intensive chemotherapy (Supplemental Table 1)

| | Overall Survival | | |
|---|---|---|---|
| **Variable** | HR | 95% CI | P |
| **Age** (>60 vs. ≤60 years) | 4,699 | 2,030-10,877 | **<0,001** |
| **ELN 2017** (intermediate vs. favorable risk) | 0,955 | 0,348-2,621 | 0,928 |
| **ELN 2017** (favorable vs. adverse risk) | 2,082 | 0,589-7,363 | 0,255 |
| **AML subtype** (de novo vs. secondary) | 0,683 | 0,216-2,157 | 0,516 |
| **TSAT** (≤51,5 vs. >51,5%) | 2,703 | 1,172-6,234 | **0,020** |
| **Response to induction** (CR vs. no CR) | 1,946 | 0,712-5,319 | 0,194 |

ELN – European Leukemia Net; TSAT – transferrin saturation; CR – complete response; HR – hazard ratio; CI – confidence interval; Bold: significant P values (p<0.05)
